# The Prp19C subunits Cwc15 and Syf2 function in TREX occupancy and transcription elongation

**DOI:** 10.1101/2023.08.03.551875

**Authors:** Laura Henke-Schulz, Rashmi Minocha, Katja Sträßer

## Abstract

The Prp19 complex (Prp19C) is conserved from yeast to human and functions in many different processes such as genome stability, splicing and transcription elongation. In the latter, Prp19C ensures TREX occupancy at transcribed genes. TREX in turn couples transcription to nuclear mRNA export by recruiting the mRNA exporter to transcribed genes and consequently to nascent mRNAs. Here, we assess the function of the nonessential Prp19C subunits Syf2 and Cwc15 in the interaction of Prp19C and TREX with the transcription machinery, Prp19C and TREX occupancy as well as transcription elongation. Whereas both proteins are important for Prp19C-TREX interaction, Syf2 is needed for full Prp19C occupancy, and Cwc15 is important for the interaction of Prp19C with RNA polymerase II and TREX occupancy. These partially overlapping functions are corroborated by a genetic interaction between *Δcwc15* and *Δsyf2*. Finally, Cwc15 also interacts genetically with the transcription elongation factor Dst1 and functions in transcription elongation. In summary, we uncover novel roles of the nonessential Prp19C components Syf2 and Cwc15 in Prp19C’s function in transcription elongation.

## INTRODUCTION

In order for gene expression to culminate in protein synthesis, a series of dynamic and interconnected processes take place. The pre-mRNA is synthesized by RNA polymerase II (RNAPII) and is processed into mature mRNA by capping of the 5’ end, splicing and 3’ end formation. Furthermore, several proteins bind to the mRNA and package it into a messenger ribonucleoprotein particle (mRNP) (1-6). These events largely occur already co-transcriptionally. Only correctly processed and packaged mRNPs are exported from the nucleus to the cytoplasm (2,5,7,8). Here, the mRNA serves as the template for protein synthesis.

A key player of mRNP assembly is the TREX complex, which couples <u>tr</u>anscription to nuclear mRNA <u>ex</u>port and is conserved from yeast to human (9). In *S. cerevisiae*, TREX is a heterononameric complex consisting of the THO complex (Tho2, Hpr1, Mft1, Thp2 and Tex1), the SR-like proteins Gbp2 and Hrb1, the DEAD-box RNA-helicase Sub2 and the export adaptor Yra1 (10). TREX is recruited to RNAPII-transcribed genes by several mechanisms. It binds to the nascent mRNA and interacts both with the S2-phosphorylated C-terminal domain (CTD) of Rpb1, the largest subunit of RNAPII, and with the Prp19 complex (Prp19C) (11-15). Prp19C, a complex best known for its function in splicing, in turn likewise interacts with RNAPII and furthermore with Mud2 (14,16,17). Mud2, a protein that also functions in splicing, is recruited to the transcription machinery by the S2-phosphorylated CTD of Rpb1 (16,18). Thus, TREX is recruited to the transcription machinery by several interactions.

As mentioned above, Prp19C is important for TREX occupancy at transcribed genes and thus for efficient transcription elongation (15,19,20). Prp19C and its associated proteins join the spliceosome during or after the dissociation of the U4 snRNP. Prp19C stabilizes the association of the U5 and U6 snRNPs by promoting RNA-RNA interaction between U5 and U6 in the activated spliceosomal B* complex, which catalyzes the first transesterification reaction and remains bound throughout the second step of splicing (17,21-23). In addition to splicing and transcription elongation, Prp19C is involved in genome stability (24) and lipid droplet formation (25). Like the TREX complex, Prp19C is conserved from yeast to human. In humans, at least three different Prp19-like complexes exist, whereas only one Prp19C is known in yeast (20). In *S. cerevisiae*, Prp19C consists of eight core subunits and 26 associated proteins (20,26). The core subunits are the four essential proteins Prp19, Cef1, Syf1 and Clf1 and the four nonessential proteins Snt309, Syf2, Isy1 and Ntc20 (19,27). In addition to these eight core proteins in *S. cerevisiae*, the human Prp19C contains additional core proteins, namely PRL1/PRLG1, AD002/HSPC148, CTNNBL1/NAP and HSP73. The yeast orthologs of PRL1/PRLG1 and AD002/HSPC148 are the Prp19C-associated proteins Prp46 and Cwc15, respectively, while the other two proteins do not have orthologs in yeast (27).

Single deletion mutants of the nonessential Prp19C subunits do not show a strong phenotype (26,28-30). Three of the four nonessential proteins, namely Isy1, Ntc20 and Syf2, have overlapping functions and form a subcomplex together with the essential subunit Syf1 (29). The triple deletion of *ISY1, NTC20* and *SYF2* is lethal, while the different double deletions are not (28,29). Double deletion of *ISY1* and *NTC20*, however, causes a severe growth defect and a high accumulation of non-spliced mRNAs (28). Deletion of *SNT309*, encoding the fourth nonessential core protein of Prp19C, leads to accumulation of non-spliced mRNAs at elevated temperatures (30). Within Prp19C, Snt309 interacts exclusively with Prp19, and the lack of Snt309 results in the destabilization of the whole complex (31). Furthermore, the associated subunit Cwc15 is involved in pre-mRNA splicing and interacts directly with the essential subunit Cef1 (26). Taken together, the nonessential proteins show no severe effects if deleted separately, but combinations of two or three deletions show more severe phenotypes. Therefore, while some of these proteins can compensate for the lack of other nonessential Prp19C components, collectively they perform essential functions.

Here, we elucidate the function of the nonessential Prp19C subunits and the associated factor Cwc15 in Prp19C and TREX occupancy at transcribed genes and in transcription elongation. We show that Syf2 plays a role in the interaction of Prp19C with TREX and in Prp19C occupancy. Cwc15 plays a role in the interaction between Prp19C and RNAPII, TREX occupancy and efficient transcription elongation. Furthermore, deletion of either *CWC15* or *SYF2* leads to a reduced interaction of Prp19C with TREX. This interaction is further decreased in a *Δcwc15 Δsyf2* double deletion strain, and *Δcwc15* and *Δsyf2* are synthetically lethal at 37°C. Interestingly, *Δcwc15* is also synthetically sick with *Δdst1* at 37°C and in the presence of 6-AU, indicating a function of Cwc15 in transcription elongation. Consistent with this conclusion, the mRNA synthesis rate is decreased in a *Δcwc15* strain. Taken together, we show that Cwc15 and Syf2 are important for the interaction between TREX and Prp19C and that Cwc15 has a role in transcription elongation.

## MATERIAL AND METHODS

### Quantification of total protein levels by Western blot

To quantify total protein levels, 5 OD_600_ units of cells grown to mid-log phase were harvested, lysed by the NaOH method and proteins precipitated by TCA (32). Briefly, the cell pellet was resuspended in 500 μl of water, 150 μl of pre-treatment solution (7.5% (v/v) β-mercaptoethanol, 1.85 M NaOH) were added, and the mixture was incubated for 20 min on ice. TCA was added to a final concentration of 10%, the solution incubated for 20 min on ice and centrifuged for 20 min at 15,000 rpm at 4°C. The supernatant was discarded, the pellet was resuspended in 90 μl of 1x SDS-loading buffer, and 10 μl 1 M Tris-base were added. Equal amounts of total protein corresponding to about 6 μl of whole cell extract were separated by SDS-PAGE. Proteins tagged with the TAP (tandem affinity purification) or HA (hemagglutinin) tag were detected with PAP or an anti-HA antibody (Sigma, Taufkirchen, Germany; R&D Systems, Minneapolis, USA) and a horseradish peroxidase (HRP)-coupled secondary antibody and CheLuminate-HRP ECL solution (Applichem) according to the manufacturer’s instructions. Western blot signals were imaged using a ChemoCam Imager (Intas) and quantified using ImageJ. Each quantification is based on at least three biologically independent replicates.

### Tandem affinity purification (TAP) of native protein complexes

Purification of native protein complexes via a TAP-tagged subunit was performed as described previously (33). Briefly, 2 l of cell culture were harvested at an OD_600_ of 3.5 and lysed with a cryo-mill (Freezer/Mill 6870D, Spex Sample Prep). Lysis buffer (50 mM Tris-HCl, pH 7.5, 100 mM NaCl, 1.5 mM MgCl_2_, 0.15% NP40, 1 mM DTT, 1.3 μg/mL pepstatin A, 0.28 μg/mL leupeptin, 170 μg/ml PMSF, 330 μg/mL benzamidine) was added to the cell powder, centrifuged (1 h, 164,700 g, 4°C) and the supernatant incubated for 1.5 h with IgG-coupled sepharose beads (GE Healthcare). The beads were washed with lysis buffer and proteins eluted by cleavage with TEV protease. The TEV eluates were incubated for 1 h with prewashed calmodulin sepharose beads (Agilent Technologies) and washed with lysis buffer containing 2 mM CaCl_2_. Proteins were eluted with buffer containing 25 mM EGTA, precipitated with 10% TCA (v/v) and separated by SDS-PAGE. Copurifying proteins were analyzed by Coomassie staining or Western blotting using antibodies against HA (Roche) or the CTD of Rpb1 (8WG16, Biolegend).

### *In vitro* binding assay of Prp19C and THO

Prp19C was purified from 12 l of culture of a *SYF1-TAP* strain until EGTA elution (EGTA-E). THO and mRNA capping enzyme, a heterotetramer composed of two Ceg1 and two Cet1 molecules that served as a negative control, were purified from an *HPR1-TAP* strain and a *CEG1-TAP* strain, respectively. Lysates of the *HPR1-TAP* and *CEG1-TAP* strains were treated with 100 μg/ml RNase A and incubated with IgG Sepharose beads. Beads were washed with 10 ml high salt lysis buffer (50 mM Tris-HCl at pH 7.5, 1 M NaCl, 1.5 mM MgCl_2_, 0.15% NP40, 1 mM DTT, 1.3 μg/ml pepstatin A, 0.28 μg/ml leupeptin, 170 μg/ml PMSF, 330 μg/ml benzamidine) in order to obtain complexes. Equal amounts of purified and concentrated Prp19C were incubated with THO or the mRNA capping complex bound to IgG-coupled Sepharose beads at 4°C for 1 h on a turning wheel. After incubation, the beads were washed with 5 ml TAP lysis buffer, the proteins were eluted by cleavage with TEV protease for 1 h at 4°C and separated by SDS-PAGE.

### Chromatin immunoprecipitation (ChIP) experiments

ChIP experiments were performed as described in (34) with some modifications. Briefly, 100 ml yeast culture were grown to an OD_600_ of 0.8 and crosslinked with 1% formaldehyde. Cells were lysed with an equal volume of glass beads and sonicated 3 times for 15 min each with intermittent cooling, resulting in chromatin fragments of 200 - 250 bp. Endogenously TAP-tagged versions of the proteins of interest were immunoprecipitated by incubation of whole cell lysate with IgG-coupled Dynabeads (tosylactivated M280, Thermo Scientific) for 2.5 h at RT. For ChIP experiments of RNAPII, 4 μl of the monoclonal antibody 8WG16 (Biolegend) were added for 1.5 h at RT followed by 1 h incubation with Protein G Dynabeads. After washing and elution, eluates as well as input samples were treated with proteinase K overnight at 65°C for the reversal of crosslinks. To determine the occupancy of each protein of interest at transcribed genes, eight paradigmatic genes were amplified from the precipitated DNA by quantitative PCR using specific primer pairs; a non-transcribed region (NTR1, 174131-174200 on chr. V) served as negative control. The occupancy of each protein was calculated as the enrichment in the immunoprecipitated (IP) over the input (Input) sample of each transcribed gene normalized to NTR1 ([E^(C_*T*_IP-C_*T*_Input)]_*NTR*_/[E^(C_*T*_IP-C_*T*_Input)]_*gene*_).

### *In vivo* transcription assay

The *in vivo* transcription assays were performed as described in (16). Shortly, transcription of two genes, the endogenous intronless *GAL10* gene and the intron-containing *ACT1* gene also under control of the GAL10 promoter and encoded on a plasmid, was examined; the RNAPIII transcript *SCR1* served as a standard. Cells were grown in media containing raffinose as a carbon source (2% w/v), and expression from the *GAL10* promotors was induced by addition of 2% (w/v) galactose for 0 - 30 min. The amount of *GAL10* and *ACT1* mRNA and *SCR1* was measured by primer extension using 5’-Cy5-labeled oligonucleotides specific for *GAL10, ACT1* and *SCR1*, respectively. The cDNA was separated on a 7 M urea 7% polyacrylamide gel and quantified using ImageJ.

### Statistical analysis

All data are presented as mean ± standard deviation (error bars) of at least three biologically independent experiments. Asterisks indicate the statistical significance (Student’s t-test; * = p-value ≤ 0.05; ** = p-value ≤ 0.01; *** = p-value ≤ 0.001).

## RESULTS

### The nonessential Prp19C subunit Snt309 is needed for the interaction of Prp19C with Mud2

Prp19C is recruited to transcribed genes by Mud2 in an RNA-independent manner (16), and the essential Prp19C subunit Clf1 most likely is important for this interaction (35). No further interactions between Prp19C components and Mud2 are known to date. We therefore investigated whether any of the nonessential Prp19C subunits are involved in the Prp19C-Mud2 interaction. To do so, we purified Prp19C from different strains expressing Syf1-TAP, each carrying a deletion of one of the nonessential Prp19C subunits, and assessed the copurification of HA-tagged Mud2 by Western blotting. To exclude any differences in the interaction of Mud2 and Prp19C due to changes in Syf1 or Mud2 total protein levels caused by one of the deletions, we determined the amount of Syf1 and Mud2 in whole cell lysates of each deletion mutant. Both Syf1 and Mud2 levels are constant in each of the deletion mutants (Supplementary Figure S1A and B). Interestingly, deletion of *ISY1* leads to an increased interaction between Prp19C and Mud2 (Figure 1A and B). In contrast, the interaction between Prp19C and Mud2 is significantly reduced when *SNT309* is deleted (Figure 1A and B). A possible explanation for this weaker interaction could be the lower integrity of Prp19C in *Δsnt309* cells (31), since a weaker association of Clf1 with Prp19C could in turn weaken the interaction of Prp19C with Mud2. Nevertheless, Snt309 does play a role in the interaction between Prp19C and Mud2.

**Figure 1.**
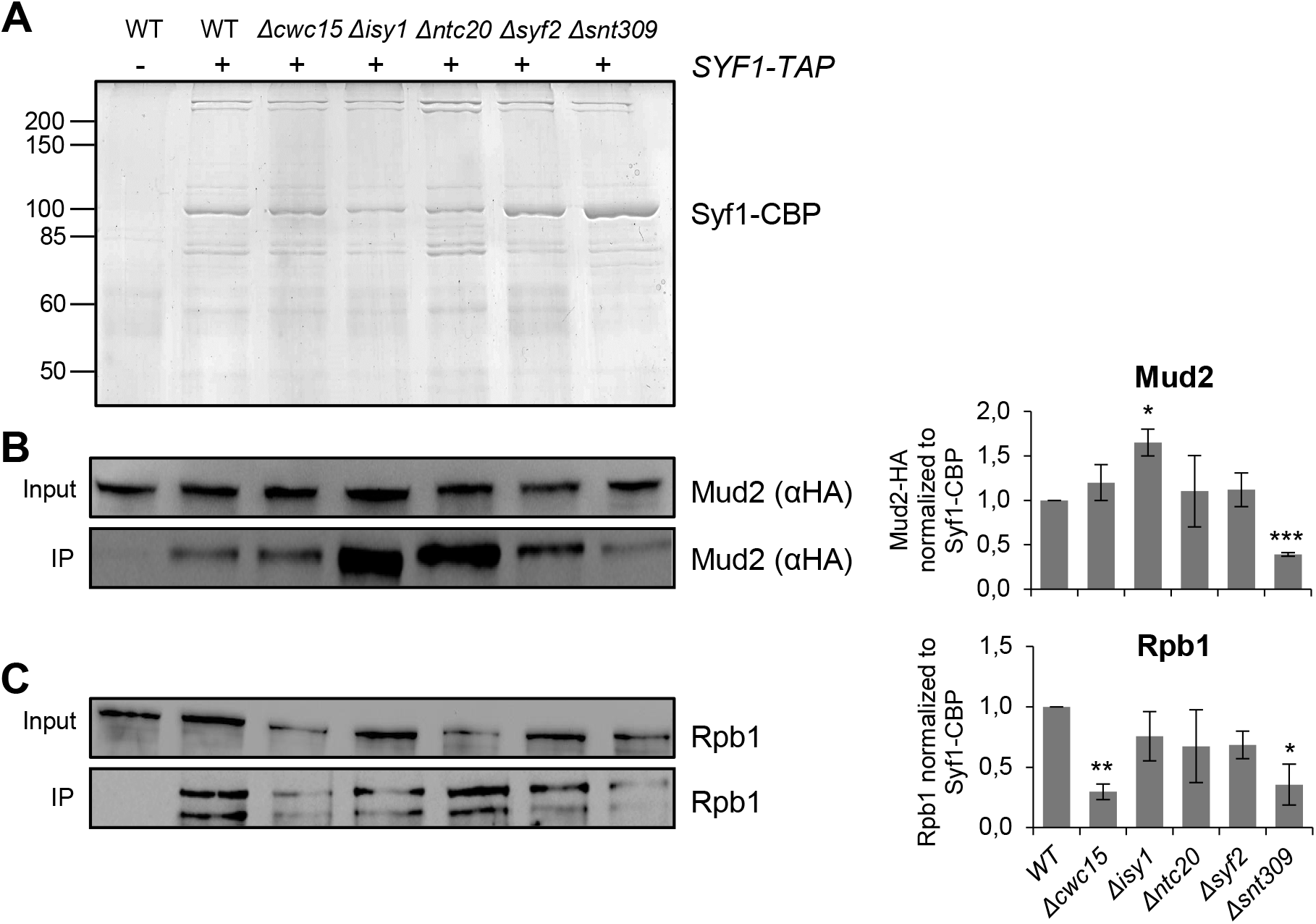
Snt309 is necessary for the interaction between Prp19C and Mud2, and Snt309 and Cwc15 are important for the interaction of Prp19C with RNAPII. (**A**) Coomassie gel of the EGTA eluates of Prp19C purified via TAP-tagged Syf1 from wild-type (WT) and the five deletion strains. The levels of copurifying HA-tagged Mud2 and Rpb1 was assessed by Western blotting with an antibody directed against HA (αHA) or the antibody 8WG16 recognizing Rpb1, respectively. A strain expressing Mud2-HA only, i.e., lacking Syf1-TAP, served as negative control. (**B and C**) Exemplary Western blot of three independent experiments using αHA (B) and 8WG16 (C) antibodies as described above (left panels) and quantification of the respective bands normalized to the essential Prp19C component Syf1 (right panels.

### The Prp19C subunits Cwc15 and Snt309 mediate the interaction between Prp19C and RNAPII

The essential Prp19C subunit Syf1 interacts – directly or indirectly – with the large subunit of RNAPII (36). Thus, we were interested whether any of the nonessential Prp19C components are important for the interaction between these two complexes. The copurification of RNAPII with Prp19C was assessed in the five deletion mutants of each of the nonessential Prp19C components (Figure 1A). To exclude any effects by reduced total RNAPII levels caused by deletion of one of these genes, we determined the total levels of the largest RNAPII subunit Rpb1 in whole cell extracts. A minor decrease of total Rpb1 levels occurs in *Δntc20* cells (Supplementary Figure 1C); however, we did not observe a significant difference in the interaction of Prp19C with Rpb1 in *Δntc20* cells (Figure 1A and C). In contrast, the interaction between Prp19C and Rpb1 and thus most likely the whole RNAPII complex decreases in *Δcwc15* and *Δsnt309* cells (Figure 1A and C). Thus, Cwc15 and Snt309 contribute to the interaction between Prp19C and RNAPII.

### Cwc15, Syf2 and Snt309 play an important role in the interaction of Prp19C and TREX

Prp19C is important for the recruitment of TREX to actively transcribed genes and interacts with TREX in an RNA-independent manner *in vivo* (36). Thus, we determined, whether Prp19C and THO interact directly using an *in vitro* binding experiment. First, we purified THO and the heterotetrameric mRNA capping enzyme, the latter of which served as a negative control, bound to IgG beads. During the purification, the lysates were treated with RNaseA to prevent any RNA-mediated interactions. In addition, the IgG beads with bound THO or capping complex were washed with high salt buffer to obtain pure complexes. Equal amounts of Prp19C purified until the EGTA elution step were incubated with THO or the mRNA capping enzyme bound to beads, respectively. Bound proteins were eluted by cleavage with TEV protease. Prp19C indeed binds directly to the THO complex (Supplementary Figure S2). In contrast, no interaction was observed between Prp19C and the mRNA capping enzyme (Supplementary Figure S2). Thus, Prp19C binds directly to the THO complex in an RNA-independent manner.

Next, we determined which of the nonessential Prp19C components are involved in this direct interaction between Prp19C and TREX. To do this, we purified TREX using a strain expressing Hpr1-TAP in deletion mutants of each of the nonessential Prp19C subunits and assessed the copurification of Prp19C by Western blotting for HA-tagged Syf1. The amount of copurified Prp19C increases when *ISY1* is deleted, indicating that the interaction between the two complexes is stronger in the absence of Isy1 (Figure 2A and B). In deletion mutants of *CWC15, SYF2* and *SNT309*, the amount of copurified Syf1 and thus the interaction of TREX with Prp19C is reduced (Figure 2A and B).

**Figure 2.**
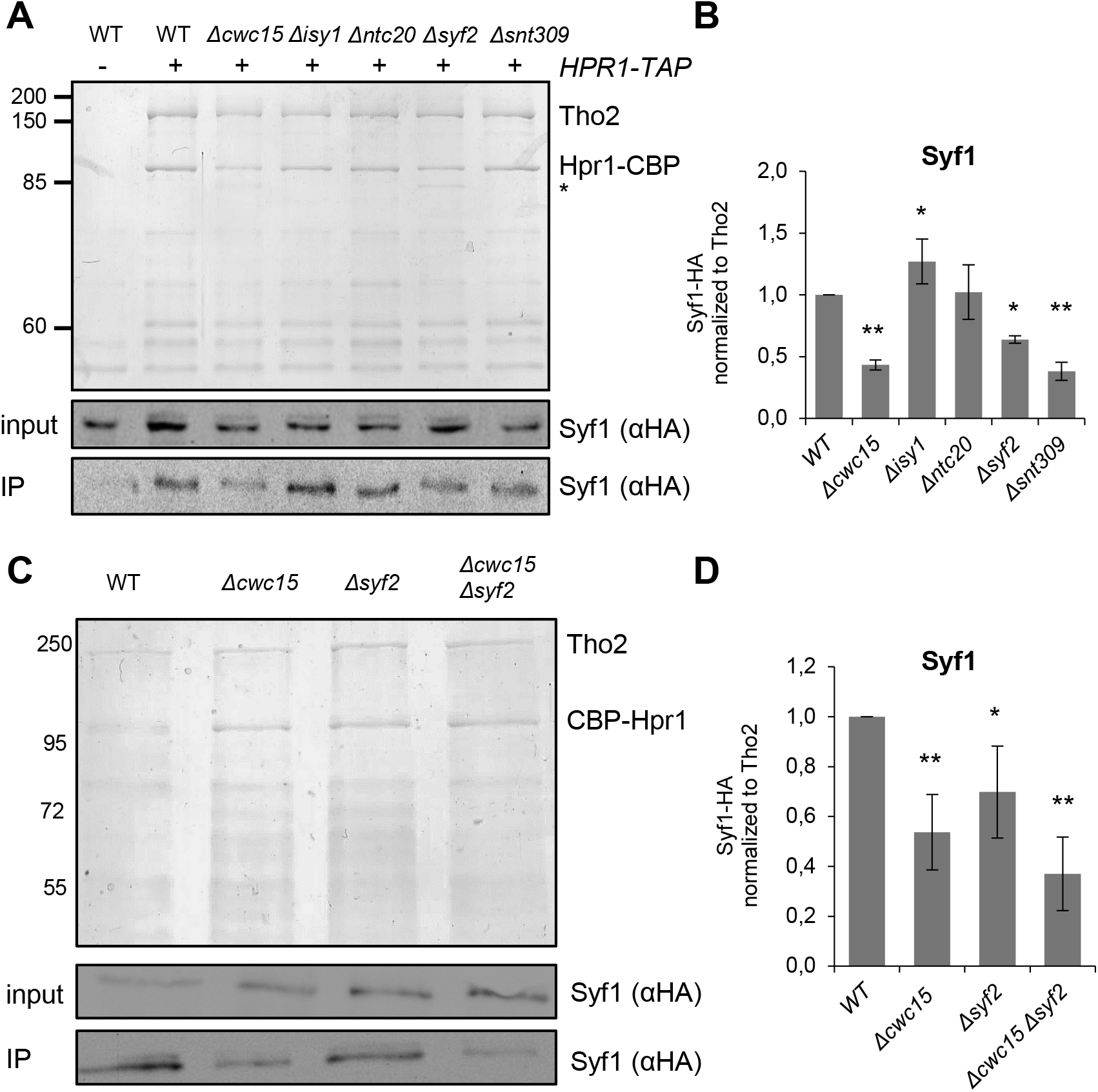
Cwc15, Syf2 and Snt309 are needed for the interaction between TREX and Prp19C. (**A**) Coomassie gel of EGTA eluates from cells expressing Hpr1-TAP and HA-tagged Syf1 in a wild-type (WT), *Δcwc15, Δisy1, Δntc20, Δsyf2* and *Δsnt309* background (upper panel). CBP, calmodulin binding protein; the faster migrating band of Hpr1-CBP in *Δcwc15* and *Δsyf2* cells was verified by mass spectrometry and is indicated by a star. Syf1-HA levels of extracts (input) and eluates (IP) were assessed by Western blotting using antibody against the HA-tag (lower panels). A strain expressing Syf1-HA only, i.e., lacking the TAP tag on Hpr1, served as negative control. (**B**) Quantification of Western blots of three independent copurifications as shown in (A). (**C**) Coomassie gel of EGTA eluates of purified, N-terminally TAP-tagged Hpr1 from wild-type, *Δcwc15, Δsyf2* and *Δcwc15Δsyf2* cells (upper panel) and Western blots assessing the corresponding Syf1-HA levels of extracts (input) and eluates (IP) (lower panels). (**D**) Quantification of Syf1 copurified with TREX of three independent experiments as shown in (C).

Interestingly, the total Hpr1-TAP levels in whole cell lysates also decrease in the absence of either Cwc15 or Syf2 as determined with an antibody directed against the protein A moiety of the C-terminal TAP tag (Supplementary Figure 3A and B). These lower Hpr1 levels coincide with a faster migrating band of Hpr1-CBP visible in the TREX eluates from *Δcwc15* and *Δsyf2* cells (Figure 2A, indicated by a star). To investigate whether the double deletion of *CWC15* and *SYF2* has additive effects on the amount of the faster migrating Hpr1-CBP band as well as the TREX-Prp19C interaction, we purified TREX from a *Δcwc15 Δsyf2* double deletion strain. Only the faster migrating band of Hpr1-CBP is visible in the EGTA eluate from this *Δcwc15 Δsyf2* strain (Supplementary Figure 3C, indicated by a star). In addition, the interaction of TREX with Prp19C is further decreased (Supplementary Figure 3C and D). Surprisingly, total levels of Hpr1-TAP are restored in the *Δcwc15 Δsyf2* strain (Supplementary Figure 3E and F). Mass spectrometric analysis of the Hpr1-CBP bands purified from the wild-type and the double deletion strain revealed peptides corresponding to full-length Hpr1 in both cases (data not shown). As Hpr1 is ubiquitylated (37,38), the size difference could be due to ubiquitylation. Detection of ubiquitylated Hpr1 in the Hpr1-TAP eluates by Western blotting with an antibody directed against ubiquitin indeed revealed strongly decreased ubiquitylation of Hpr1 in the double mutant compared to wild-type cells (Supplementary Figure 4A and B). Hpr1 is ubiquitinylated at its C-terminus by the E3 ubiquitin ligase Rsp5, destabilizing Hpr1 at 37°C (38). Indeed, Hpr1 is degraded more quickly at 37°C than at 30°C (Supplementary Figure 4C and (38)). Deletion of *CWC15* and *SYF2* increases the half-life of Hpr1-TAP at 37°C, consistent with its lower ubiquitylation level (Supplementary Figure 4C). Thus, the C-terminal TAP tag impairs ubiquitylation of Hpr1 when Cwc15 and/or Syf2 is lacking.

**Figure 3.**
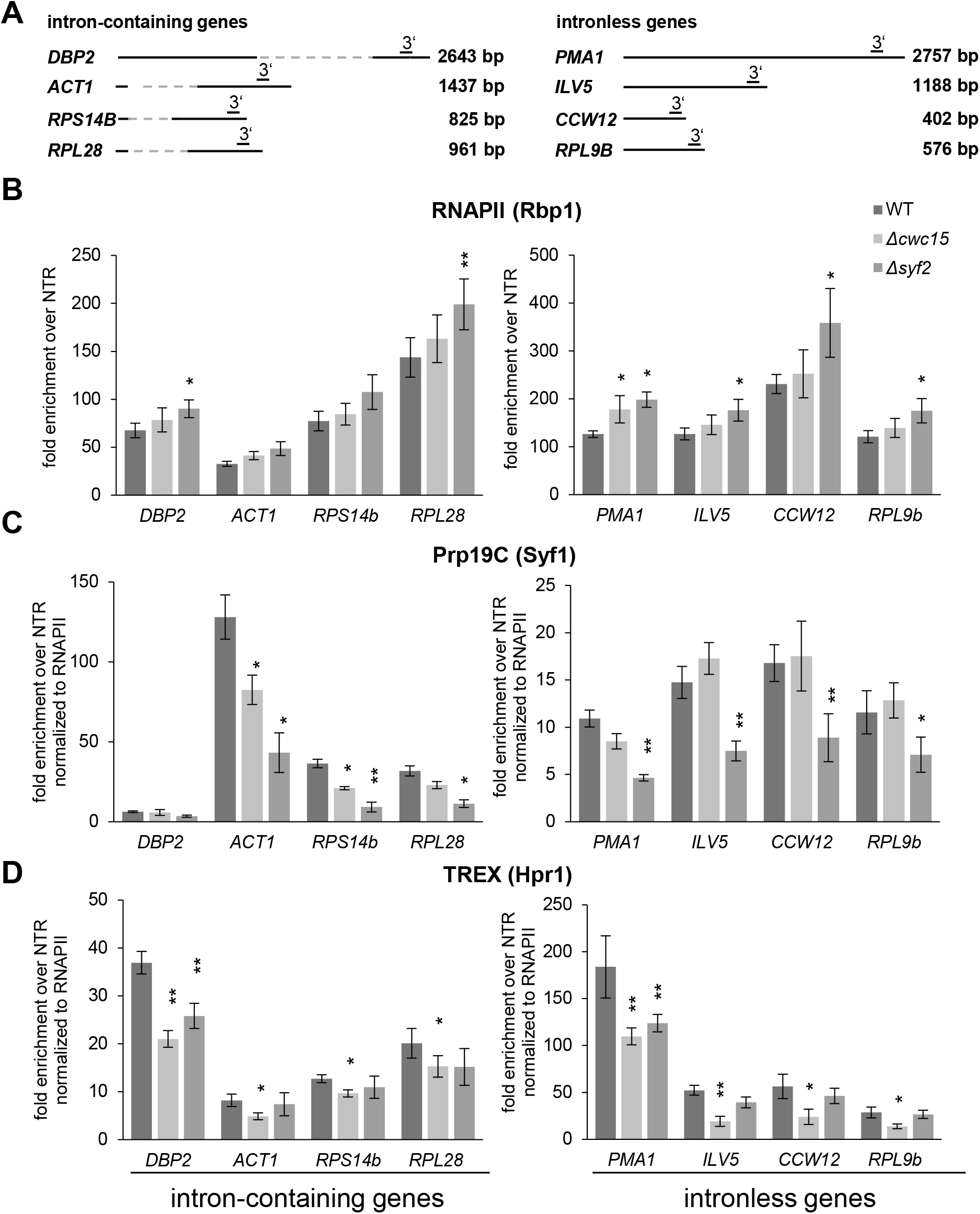
Syf2 functions in Prp19C occupancy, and Cwc15 functions in TREX occupancy. (**A**) Scheme of four exemplary intron-containing genes, *DBP2, ACT1, RPS14B* and *RPL28*, and four exemplary intronless genes, *PMA1, ILV5, CCW12* and *RPL9B* used for chromatin immunoprecipitation (ChIP) experiments. Open reading frames (ORFs) are represented by solid lines and introns by hatched lines. The bars above the genes indicate the positions of the primer pairs used for analysis by quantitative PCR. (**B**) RNAPII occupancy increases in *Δsyf2* cells both at intron-containing and intronless genes. The occupancy of the RNAPII component Rpb1 was assessed in wild-type (WT), *Δcwc15* and *Δsyf2* mutant cells by ChIP experiments at the four intron-containing (left panel) and the four intronless (right panel) genes depicted in (A). (**C**) Prp19C occupancy decreases in *Δsyf2* mutant cells at intron-containing genes. The occupancy of Prp19C was assessed by ChIP of TAP-tagged Syf1 normalized to the occupancy of RNAPII. (**D**) TREX occupancy decreases in *Δcwc15* cells. The occupancy of TREX was assessed by ChIP of TAP-tagged Hpr1 normalized to the occupancy of RNAPII.

**Figure 4.**
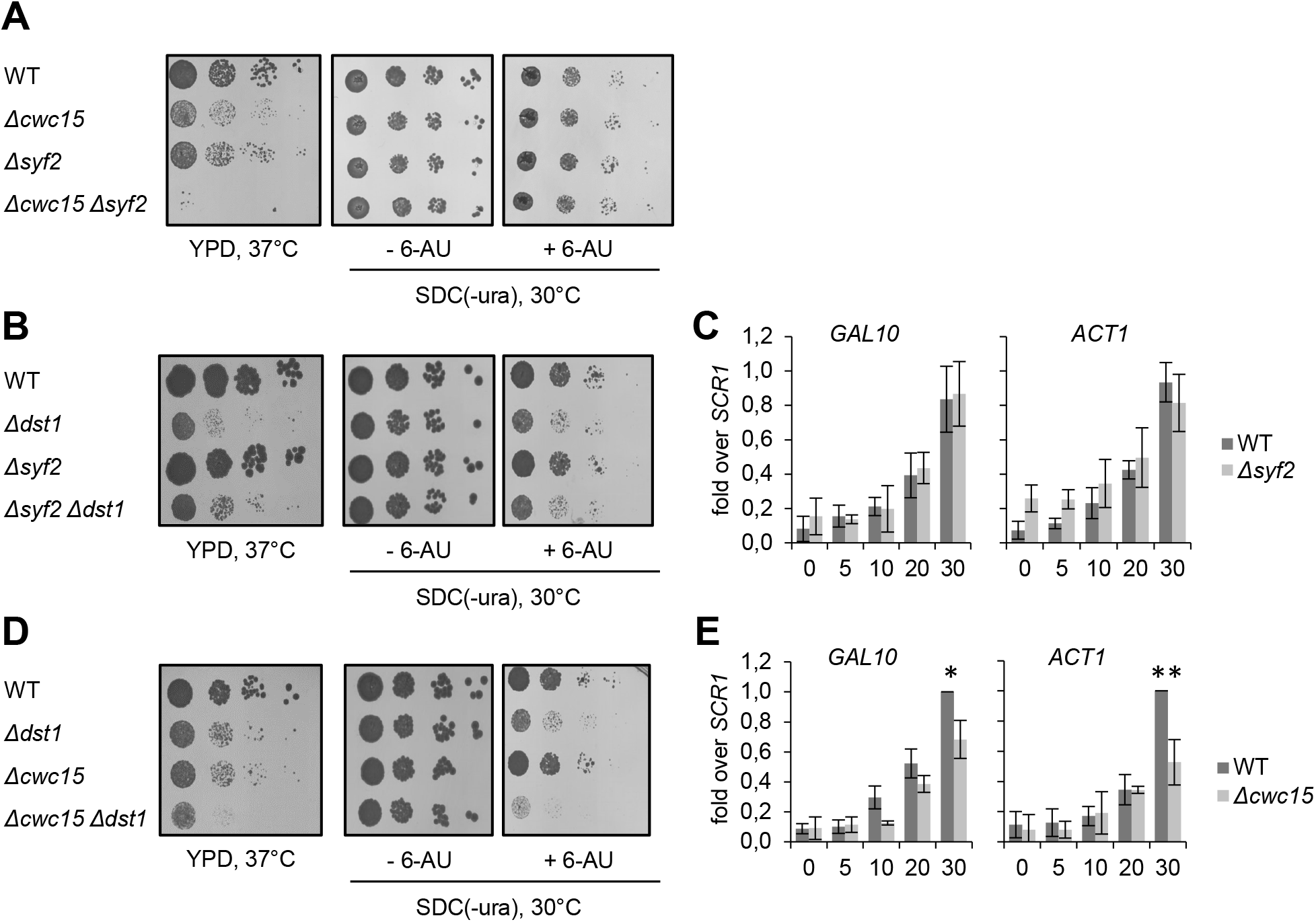
Cwc15 activity is needed for full transcriptional activity. (**A**) *CWC15* and *SYF2* interact genetically. *Δcwc15* and *Δsyf2* are synthetically lethal at 37°C. Ten-fold serial dilutions wild-type (WT), *Δcwc15, Δsyf2* and *Δcwc15 Δsyf2* cells were spotted on YPD plates and incubated for 3 days at 37°C (left panel) and on SDC(-ura) plates containing solvent (−6-AU) or 75 μg/ml 6-AU (+6-AU) and incubated for 2-3 days at 30°C (right panel). (**B**) Deletion of *SYF2* does not cause a temperature-sensitive phenotype and does not interact genetically with *Δdst1*. Dot spots of WT, *Δsyf2, Δdst1* and *Δdst1 Δsyf2* cells as described in (A). (**C**) Syf2 is not needed mRNA synthesis *in vivo*. Expression of the endogenous, intronless *GAL10* gene and the plasmid-encoded, intron-containing *ACT1* gene driven by the *GAL10* promoter was induced for 5, 10, 20 and 30 min by addition of galactose. Total RNA was extracted and the amount of *GAL10* and *ACT1* mRNA was determined by primer extension. The amount of *GAL10* and *ACT1* mRNA at the different time points of three independent experiments in wild-type (WT) and *Δsyf2* cells was quantified and normalized to the levels of the RNAPIII transcript *SCR1*. (**D**) Deletion of *CWC15* causes a temperature-sensitive phenotype and is synthetically sick with *Δdst1* both at 37°C and on 6-AU plates. Dot spots of wild-type (WT), *Δcwc15, Δdst1* and *Δdst1 Δcwc15* cells grown under the conditions described for (A). (**E**) Cwc15 is required for efficient mRNA synthesis *in vivo*. Experiment as in (C).

In order to assess the effect of Cwc15 and Syf2 on the interaction of TREX with Prp19C independent of changes in Hpr1 ubiquitylation, we tagged Hpr1 on its N-terminus. No faster migrating band is visible in the EGTA eluates of TREX purifications from *Δcwc15, Δsyf2* and *Δcwc15 Δsyf2* cells expressing TAP-Hpr1 (Figure 2C). Consistently, the total levels of TAP-Hpr1 in whole cell lysates of *Δcwc15* and *Δsyf2* cells are not reduced (Supplementary Figure 3G and H). Importantly, as for the C-terminally TAP-tagged Hpr1 (Supplementary Figure 3C and D), less Prp19C copurifies with TREX in *Δcwc15* and *Δsyf2* cells when purified via the N-terminally TAP-tagged Hpr1, and even less Prp19C copurifies with the N-terminally TAP-tagged Hpr1 in the *Δcwc15 Δsyf2* double deletion strain (Figure 2C and D). Thus, Cwc15 and Syf2 are important for the interaction of Prp19C and TREX.

### Cwc15 and Syf2 function in the recruitment of Prp19C and TREX to transcribed genes

As Cwc15 and Syf2 are needed for the interaction of Prp19C with TREX and RNAPII, we tested whether these two Prp19C components are also important for the occupancy of Prp19C and TREX at transcribed genes. As the recruitment of RNA-binding protein complexes to the site of transcription is at least partially dependent on RNA and thus ongoing transcription, we first determined the occupancy of RNAPII at four exemplary, highly transcribed intron-containing (*DBP2, ACT1, RPS14, RPL28*) and four intronless genes (*PMA1, ILV5, CCW12, RPL9B*) by chromatin immunoprecipitation (ChIP) experiments (Figure 3A). In *Δcwc15* cells, the occupancy of RNAPII is unchanged compared to wild-type cells (Figure 3B). Prp19C occupancy decreases at half of the selected intron-containing genes, whereas it does not change at intronless genes in *Δcwc15* cells (Figure 3C and Supplementary Figure S5A). TREX occupancy decreases at all tested intron-containing and intronless genes in *Δcwc15* cells (Figure 3D and Supplementary Figure S5B). This is consistent with our observation that Cwc15 is needed for a stable Prp19C-TREX interaction and the requirement of Prp19C for TREX occupancy (Figure 2 and (36)). In *Δsyf2* cells, in contrast, RNAPII occupancy increases at both intron-containing and intronless genes (Figure 3B). Prp19C occupancy decreases, whereas TREX occupancy is not affected in *Δsyf2* cells (Figure 3C and D). Taken together, Syf2 is needed for Prp19C occupancy, and Cwc15 is needed for TREX occupancy.

### Cwc15 is important for efficient transcription *in vivo*

Since deletion of *SYF2* or *CWC15* leads to similar phenotypes such as reduced TREX-Prp19C interaction, we also tested for a genetic interaction between these two genes as a genetic interaction indicates a function in the same cellular process. Interestingly, deletion of both *SYF2* and *CWC15* causes a strong additive growth defect at 37°C compared to the single deletions (Figure 4A), indeed suggesting overlapping functions of Syf2 and Cwc15. However, *SYF2* and *CWC15* do not interact genetically in the presence of 6-azauracil (6-AU) (Figure 4A). 6-AU leads to decreased intracellular GTP and UTP levels, thereby impeding RNA synthesis. Consequently, sensitivity to 6-AU indicates a function of the deleted gene in transcription elongation. Taken together, Syf2 and Cwc15 likely have overlapping functions, but this overlap appears not to be in transcription elongation.

Nevertheless, Syf2 and Cwc15 are both needed for Prp19C and TREX occupancy, and these two complexes function in transcription elongation (Figure 3 and (36)). We, therefore, further investigated the potential function of either Syf2 or Cwc15 in transcription elongation. As a first indication, we tested for a genetic interaction between *SYF2* and *DST1*, the gene encoding the transcription elongation factor TFIIS (39). However, deletion of *SYF2* does not lead to a growth defect at 37°C, and the double deletion of *SYF2* and *DST1* does not show any additive growth defect (Figure 4B, left panel). Similarly, *Δsyf2* cells are not sensitive to 6-AU and do not interact with *Δdst1* in the presence of 6-AU (Figure 4B, right panel). As an independent line of evidence, we assessed mRNA synthesis in an *in vivo* transcription assay using two reporter genes: the endogenous, intronless *GAL10* gene and the intron-containing *ACT1* gene encoded on a plasmid, likewise under control of the *GAL10* promotor. Expression from these two *GAL10* promoters was induced by addition of galactose for 5, 10, 20 and 30 minutes, respectively, and the amount of synthesized *GAL10* and *ACT1* mRNA was quantified by primer extension. However, the amount of synthesized *GAL10* and *ACT1* mRNA in *Δsyf2* cells is similar to the respective mRNA levels in wild-type cells, also indicating that Syf2 function is dispensable for efficient mRNA synthesis (Figure 4C).

In contrast to *Δsyf2* cells, *Δcwc15* cells have a growth defect comparable to that of *Δdst1* cells at 37°C (Figure 4D, left panel). Furthermore, the *Δcwc15 Δdst1* double deletion strain has an additive growth defect at 37°C, i.e., *CWC15* and *DST1* interact genetically (Figure 4D, left panel), indicating a function of Cwc15 in transcription elongation. Growth of *Δcwc15* cells on 6-AU containing plates is not impaired compared to wild-type cells; however, deletion of *CWC15* exacerbates the growth defect of *Δdst1* cells in the presence of 6-AU (Figure 4D, right panel), again indicating a role of Cwc15 in transcription elongation. To substantiate this conclusion, we performed *in vivo* transcription assays with the two above-mentioned reporter genes. Consistent with the afore-mentioned data, both *GAL10* and *ACT1* mRNA levels are significantly lower in *Δcwc15* cells compared to wild-type cells (Figure 4E). Consequently, Cwc15 is needed for efficient mRNA synthesis or stability of intron-containing and intronless mRNA *in vivo*.

Taken together, our data show that the two nonessential Prp19C subunits Cwc15 and Syf2 play similar but different roles in the interaction of Prp19C with TREX, Mud2 and RNAPII. Importantly, Cwc15 is needed for efficient transcription elongation *in vivo*.

## DISCUSSION

The TREX complex functions in transcription elongation and mRNA export (10). TREX is recruited to the transcription machinery by several mechanisms: i) by direct interaction with the S2-phosphorylated CTD, ii) by the nascent RNA and iii) by the Prp19 complex (11,14). Prp19C in turn is recruited to transcribed genes i) by Mud2, which also interacts with the S2-phosphorylated CTD, and ii) via direct interaction with RNAPII, mediated by the C-terminal domain of its subunit Syf1 (14,16).

The aim of this study was to elucidate the role of the nonessential Prp19C components in the interaction with RNAPII, Mud2 and TREX, in Prp19C and TREX occupancy and in transcription elongation. Our data show that the Prp19C subunit Snt309 is important for the interaction between Prp19C and Mud2, RNAPII as well as TREX (Figure 1 and 2), which is reflected by the growth impairment caused by deletion of *SNT309*. Snt309 is required for the stable association of Cef1, an essential Prp19C core component, with Prp19C, and deletion of *SNT309* thus destabilizes the complex (40). In addition, the strong growth defect of *Δsnt309* cells renders the analysis of Snt309’s function difficult due to secondary effects.

Cwc15 is needed for the interaction between Prp19C and RNAPII as well as TREX (Figures 1 and 2). Consistently, TREX occupancy is reduced in *Δcwc15* cells (Figure 3) and transcriptional activity of two reporter genes is reduced in *Δcwc15* compared to wild-type cells (Figure 4E). This is also consistent with the observed genetic interaction between *Δcwc15* and *Δdst1* on 6-AU plates and at 37°C (Figure 4D). In this respect, the phenotypes of *Δcwc15* cells are similar to those of *Δmud2* cells (16). This is interesting as Cwc15 is known to be a loosely attached protein of Prp19C, and for *S. cerevisiae* it is not considered a core component of Prp19C (26,41).

The nonessential Prp19C component Syf2 in turn stabilizes the Prp19C-TREX interaction and is needed for full Prp19C occupancy at transcribed genes (Figures 2 and 3). However, Syf2 forms a Prp19C subcomplex together with Syf1, Isy1 and Ntc20, and the reduced Prp19C occupancy in *Δsyf2* cells could thus be an indirect effect caused by destabilization of this subcomplex and, consequently, of the interaction of Syf1 with the transcription machinery. Nevertheless, Cwc15 and Syf2 are both needed for Prp19C-TREX interaction, and this interaction is further reduced when both proteins are missing (Figure 2C and Supplementary Figure 3C and D). This is reflected by their genetic interaction: *Δcwc15* and *Δsyf2* are synthetic lethal at 37°C (Figure 4A).

As determined by the recently published cryo-electron microscopy (cryo-EM) structures of activated spliceosomal complexes B and C, Syf2 helps to bridge Syf1 to the U2-U6 snRNA helix II during splicing, anchoring Prp19C to the active site through its component Cef1 (42,43). Syf2 thus helps to stabilize the catalytic core, and its deletion could disrupt this stable conformation which may in turn render the whole Prp19 complex unstable. Cwc15, which was earlier thought to be a spliceosome-associated protein, has been regarded as a core component of the spliceosomal machinery in recent cryo-EM studies. It is in direct contact with all three snRNA elements, U4, U5 and U6, and interacts simultaneously with multiple components and other NTC-related proteins at the catalytic center of the spliceosome (44). Together with Prp45 and a C-terminal fragment of Cef1, Cwc15 acts as a rope-like structure providing flexible tethering to seal together the interactions between different components of Prp19C as well as the spliceosome (45). These structural data support our findings that the roles of Syf2 and Cwc15 in stabilizing Prp19C indeed overlap.

The total levels of C-terminally TAP-tagged Hpr1 decrease in *Δcwc15* and *Δsyf2* cells (Supplementary Figure 3, A and B), and a faster migrating band appears in the TREX purification (Supplementary Figure 3, C and D). Interestingly, total Hpr1-TAP levels are restored in *Δcwc15 Δsyf2* double deletion strain (Supplementary Figure 3, E and F), and only the faster migrating band is visible (Supplementary Figure 3, C and D). As determined by mass spectrometric analysis, both the slower and the faster migrating band of Hpr1-CBP from wild-type and *Δcwc15 Δsyf2* cells, respectively, contain full-length protein (data not shown). Hpr1 (Hpr1-HA) is ubiquitylated by the ubiquitin ligase Rsp5, degraded by the proteasome and thus more unstable at 37°C (38). However, ubiquitylated Hpr1 also recruits the mRNA exporter Mex67 to transcribed genes, and Rps5 is essential for nuclear mRNA export (46,47). As Hpr1-TAP is less ubiquitylated in the *Δcwc15 Δsyf2* strain (Supplementary Figure 4, A), the faster migrating band most likely corresponds to non-ubiquitylated Hpr1. Consistently, this non-ubiquitylated Hpr1-TAP is more stable at 37°C in *Δcwc15 Δsyf2* than in wild-type cells (Supplementary Figure 4C), a similar effect to the one observed in *Δrsp5* cells (38). Thus, the combination of a C-terminal TAP tag on Hpr1 with deletion of *CWC15* and/or *SYF2* interferes with ubiquitylation of Hpr1.

*Δcwc15, Δsyf2* and *Δcwc15 Δsyf2* cells do not show a nuclear mRNA export defect at 30°C or 37°C (data not shown). This is comparable to deletion of the C-terminus of Syf1 in *syf1-37* cells or deletion of *MUD2*, both of which also cause a decrease in TREX and Prp19C occupancy, but no nuclear mRNA export defect (14,16). Most likely, the decrease in transcription elongation and thus in mRNA synthesis caused by the mutations alleviates the possibly existing delay in mRNA export.

In summary, Cwc15 and Syf2 mediate the interaction of TREX and Prp19C with each other as well as with the transcription machinery. In addition, Cwc15 is needed for efficient transcription elongation. Notably, this is the first time a specific function of these nonessential Prp19C subunits beyond their role in splicing has been determined.

## SUPPLEMENTARY DATA

Supplementary Data are available at NAR online.

## ACKNOWLEDGEMENT

We thank Günter Lochnit for the analysis of the Hpr1-CBP band purified from wild-type and *Δcwc15 Δsyf2* cells. We thank Sittinan Chanarat for strain *SYF1-TAP* (BY4741), Susanne Röther for strain *HPR1-TAP* (BY4741) and Eleni Karakasili for strain *Δdst1* (W303). We thank Vera Bettenworth for critical reading of the manuscript.

## FUNDING

This work was supported by the EU [ERC Consolidator grant “mRNP-PackArt” to K.S.].

## CONFLICT OF INTEREST

None declared.

## Notes

### Competing Interest Statement

The authors have declared no competing interest.

