## Supplemental Data for "The Prp19C subunits Cwc15 and Syf2 function in TREX occupancy and transcription elongation"

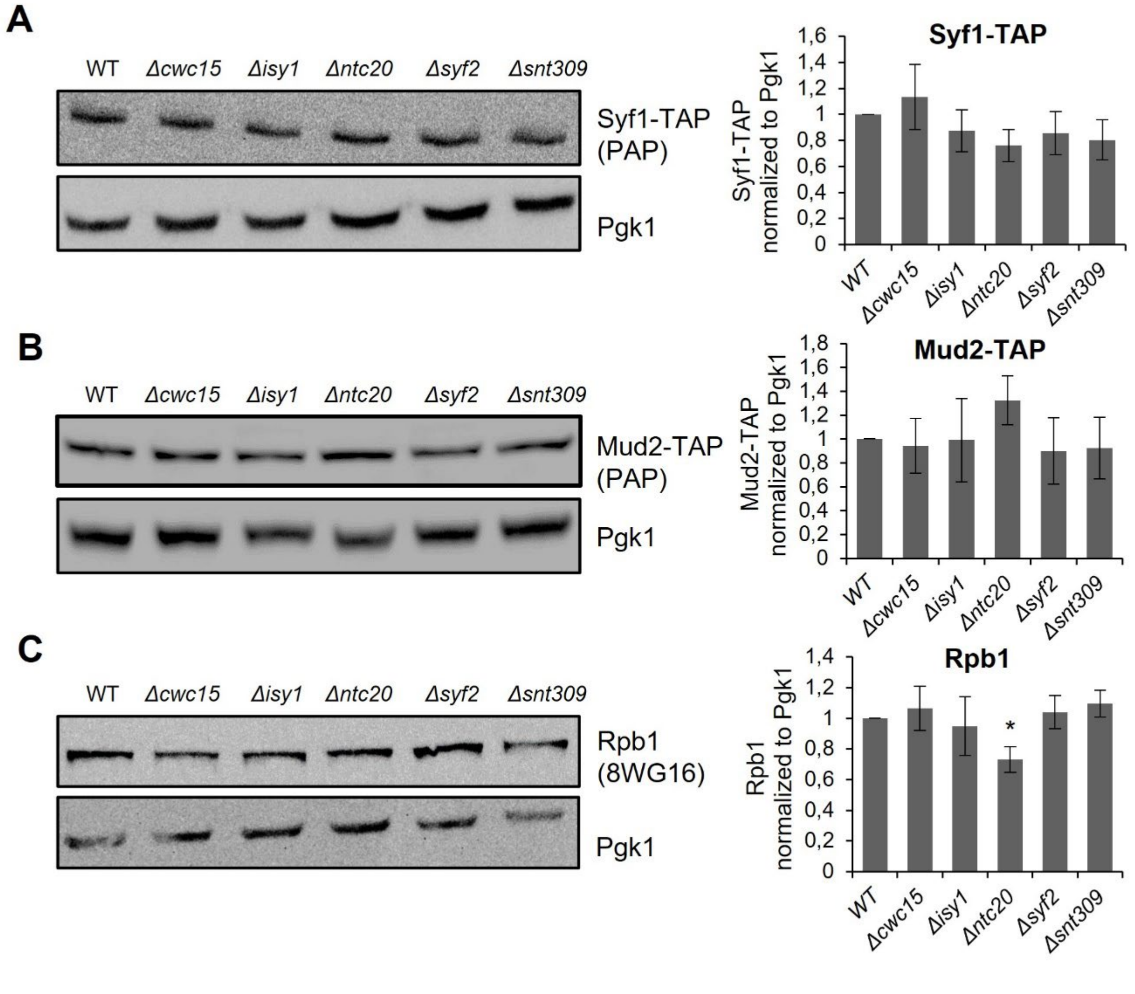


**Supplementary Figure 1.** Total levels of Syf1, Mud2 and Rpb1 do not change in deletion mutants of genes encoding the nonessential Prp19C components Cwc15, Isy1, Ntc20, Syf2 and Snt309. (**A**) Western blots (left panel) and quantification of three independent experiments (right panel) of total Syf1 levels in whole cell lysates of deletion mutants of the four nonessential Prp19C components. The amount of Syf1-TAP was quantified with an antibody directed against the protein A moiety of the TAP tag (PAP) and normalized to the amount of Pgk1. Values for wild-type (WT) cells were set to 1. (**B**) Western blots (left panel) and quantification (right panel) as in (A) except that the total levels of Mud2-TAP were determined. (**C**) Western blots (left panel) and quantification (right panel) as in (A) except that the total levels of Rpb1 were determined with antibody 8WG16. Total Rpb1 levels slightly decrease in the *Δntc20* mutant.

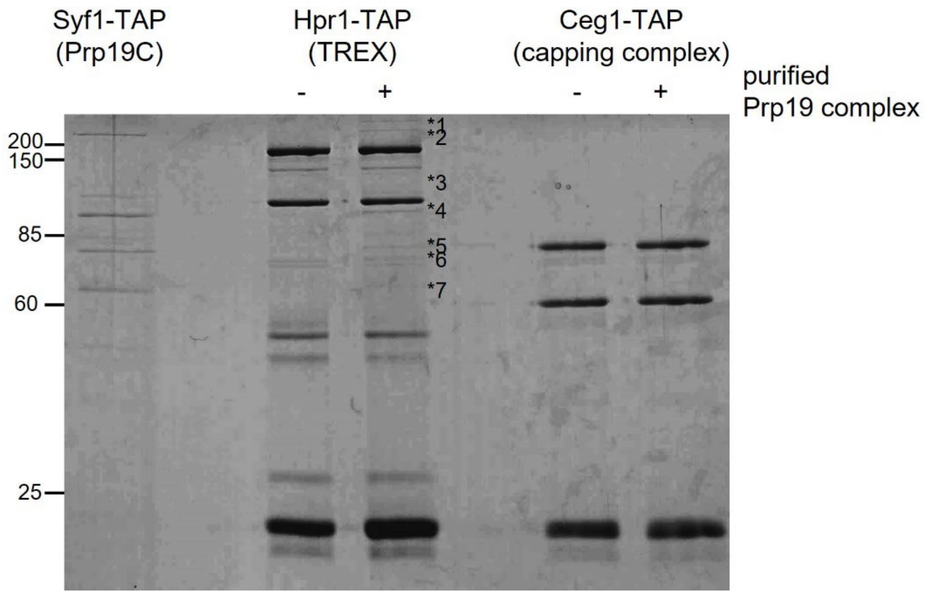


**Supplementary Figure 2.** Prp19C binds directly to the THO complex in an RNA-independent manner. Equal amounts of Prp19C were purified until the EGTA eluate from a strain expressing Syf1-TAP and incubated with purified THO and mRNA capping enzyme, a heterotetramer composed of two molecules of Ceg1 and Cet1 each that served as negative control, bound to IgG-coupled Sepharose beads. Beads were washed and the bound proteins were eluted by cleavage with TEV protease. Prp19C subunits that interact with the THO complex were identified by mass spectrometry: 1: Prp8, 2: Brr2, 3: Snu114, 4: Syf1, 5: Cef1, 6: Syf3, 7: Prp19.

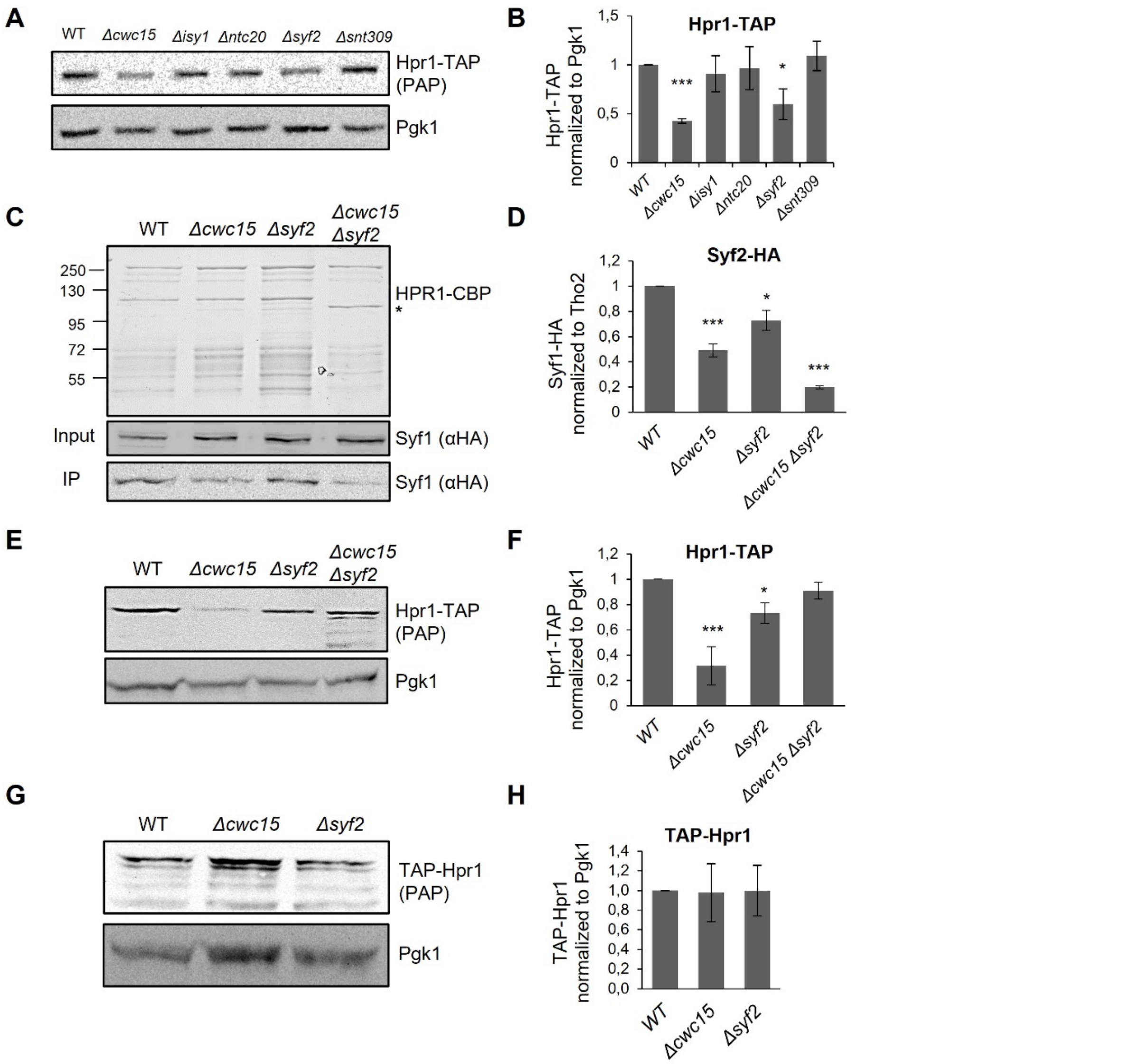


**Supplementary Figure 3.** Deletion of *CWC15* or *SYF2* causes a decrease in total Hpr1-TAP but not TAP-Hpr1 levels. (**A**) Western blots of Hpr1-TAP using an antibody against the protein A moiety of the TAP tag (PAP) in the five deletion mutants of nonessential Prp19C components compared to wild-type (WT) cells. Pgk1 levels served as a loading control. (**B**) Quantification of three independent experiments as depicted in (A). (**C and D**) Deletion of *CWC15* and/or *SYF2* decreases the interaction between TREX and Prp19C. (**C**) EGTA eluates of TREX purifications using Hpr1-TAP from strains expressing HA-tagged Syf1 in wild-type (WT), *Δcwc15*, *Δsyf2* and *Δcwc15 Δsyf2* cells. The band corresponding to Hpr1-CBP in wild-type cells is marked Hpr1-CBP, the band corresponding to faster migrating Hpr1-CBP is indicated by an asterisk (*). Syf1-HA levels of extracts (Input) and eluates (IP) were assessed by Western blotting using antibody against the HA tag. (**D**) Quantification of three independent purifications as shown in (C). (**E and F**) Double deletion of *CWC15* and *SYF2* restores total Hpr1-TAP levels. (**E**) Western blots of whole cell lysates to determine the total levels of Hpr1-TAP in wild-type (WT), *Δcwc15*, *Δsyf2* and *Δcwc15 Δsyf2* cells. Pgk1 levels served as a loading control. (**F**) Quantification of three independent experiments as shown in (E). (**G and H**) Total TAP-Hpr1 levels are unaffected in *Δcwc15* and *Δsyf2* cells. (**G**) Western blots of whole cell lysate to determine the total levels of N-terminally TAP-tagged Hpr1 in wild-type (WT), *Δcwc15* and *Δsyf2* cells. Pgk1 levels served as a loading control. (**H**) Quantification of total TAP-Hpr1 levels normalized to Pgk1 of three independent experiments.

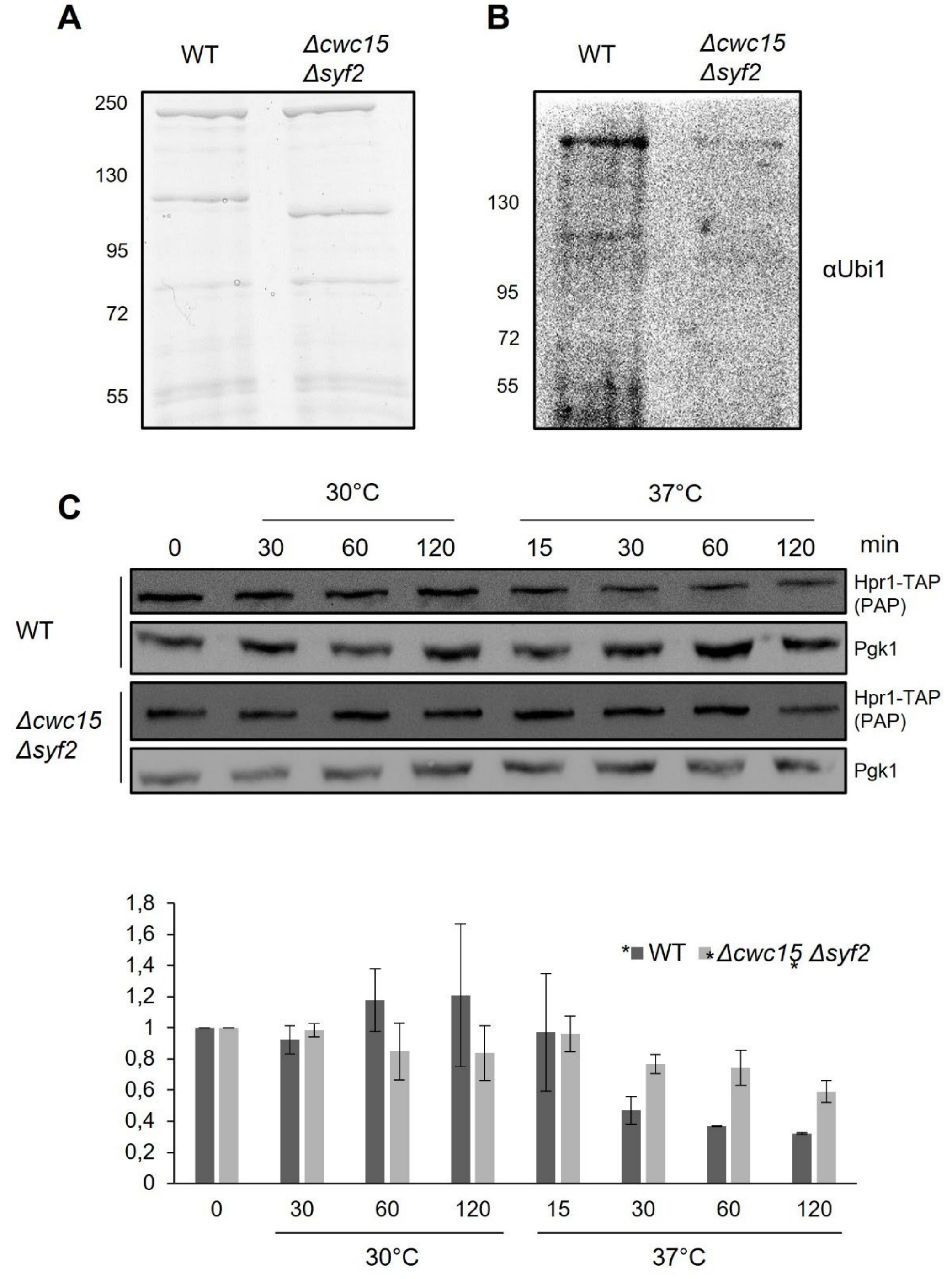


**Supplementary Figure 4**. Double deletion of *CWC15* and *SYF2* causes lower ubiquitylation levels and higher stability of Hpr1-TAP. (**A**) Coomassie gel of EGTA eluates of TAP purifications of C-terminally TAP-tagged Hpr1 from wild-type and *Δcwc15Δsyf2* cells. (**B**) Western blots of the eluates shown in (A) using an antibody against ubiquitin (αUBI1) (**C**) Hpr1-TAP is more stable in the *Δcwc15Δsyf2* double deletion mutant than in wild-type cells at 37°C. Cells were grown in YPD medium at 30°C and shifted to 37°C at an $OD_{600}$ of 0.6, and samples were taken at the time points indicated. Upper panel: Western blots with antibodies directed against protein A moiety of the TAP tag to detect Hpr1-TAP (PAP) or against Pgk1, which served as loading control. Lower panel: Relative amount of Hpr1-TAP normalized to the amount of Pgk1 at t = 0 min, based on quantification of three independent experiments as depicted in the upper panel.

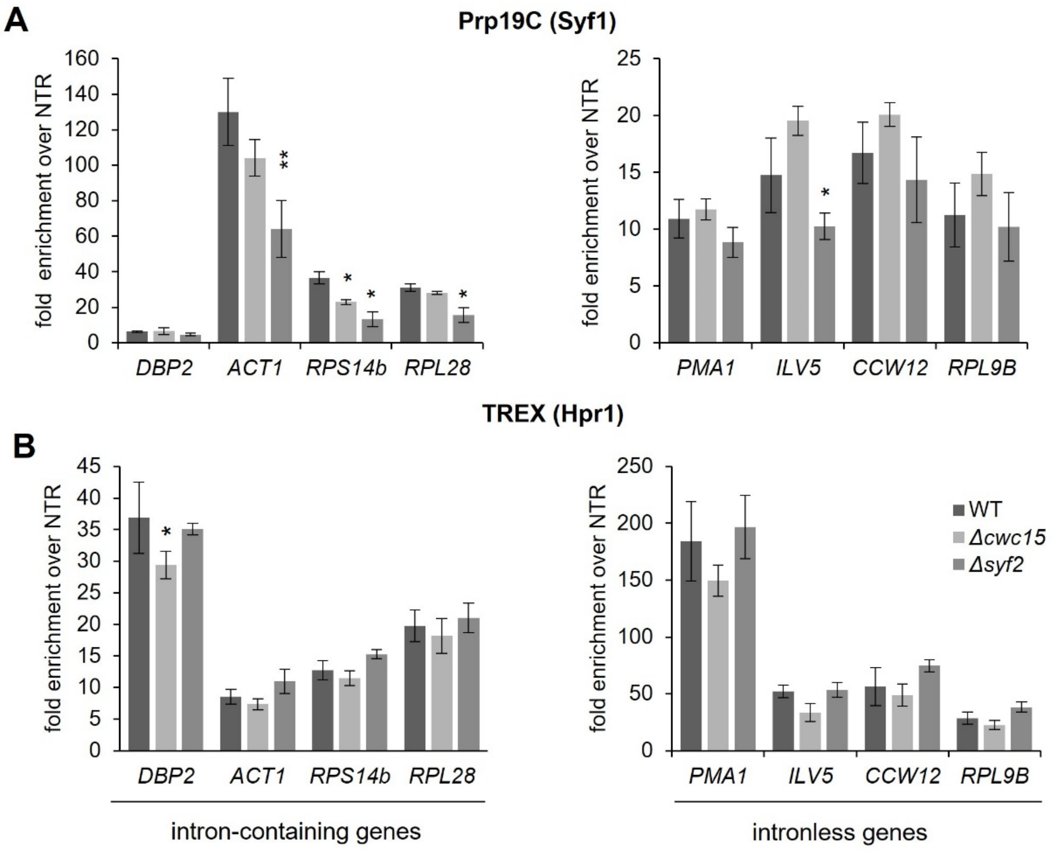


**Supplementary Figure 5.** Prp19C occupancy is reduced in *Δsyf2* cells and TREX occupancy is reduced in *Δcwc15* cells. The occupancy of Syf1 (**A**) and Hpr1 (**B**) was assessed in wild-type cells (WT), *Δcwc15* and *Δsyf2* deletion mutants using chromatin immunoprecipitation (ChIP) at both intron-containing (left panels) and intronless genes (right panels). ChIPs were quantified by Real Time PCR using primer pairs annealing to the 3' end of the respective genes.

### Supplementary Table S1. Yeast strains

| Strain | Strain Background | Genotype | Reference |
|---|---|---|---|
| wild-type | BY4741 | *MATa; his3Δ1; leu2Δ0; met15Δ0; ura3Δ0* | (1) |
| wild-type | W303 | *MATa; ade2-1; his3-11, 15; ura3-1; leu2-3, 112; trp1-1; can1-100; rad5-535* | (2) |
| *MUD2-HA* | BY4741 | *MATa; ura3Δ0; leu2Δ0; his3Δ1; met15Δ0; MUD2-HA::HIS3* | this study |
| *SYF1-TAP MUD2-HA* | BY4741 | *MATa; ura3Δ0; leu2Δ0; his3Δ1; met15Δ0; SYF1-TAP::URA, MUD2-HA::HIS* | this study |
| *SYF1-TAP MUD2-HA Δcwc15* | BY4741 | *MATa; ura3Δ0; leu2Δ0; his3Δ1; met15Δ0; SYF1-TAP::URA3; MUD2-HA::HIS3; Δcwc15::kanMX4* | this study |
| *SYF1-TAP MUD2-HA Δisy1* | BY4741 | *MATa; ura3Δ0; leu2Δ0; his3Δ1; met15Δ0; SYF1-TAP::URA3; MUD2-HA::HIS3; Δisy1::kanMX4* | this study |
| *SYF1-TAP MUD2-HA Δntc20* | BY4741 | *MATa; ura3Δ0; leu2Δ0; his3Δ1; met15Δ0; MUD2-HA::HIS3; SYF1-TAP::URA3; Δntc20::kanMX4* | this study |
| *SYF1-TAP MUD2-HA Δsyf2* | BY4741 | *MATa; ura3Δ0; leu2Δ0; his3Δ1; met15Δ0; MUD2-HA::HIS3; SYF1-TAP::URA3; Δsyf2::kanMX4* | this study |
| *SYF1-TAP MUD2-HA Δsnt309* | BY4741 | *MATa; ura3Δ0; leu2Δ0; his3Δ1; met15Δ0; MUD2-HA::HIS3; SYF1-TAP::URA3; Δsnt309::kanMX4;* | this study |
| *SYF1-HA* | BY4741 | *MATa; ura3Δ0; leu2Δ0; his3Δ1; met15Δ0; SYF1-HA::HIS3* | this study |
| *HPR1-TAP SYF1-HA* | BY4741 | *MATa; ura3Δ0; leu2Δ0; his3Δ1; met15Δ0; SYF1-HA::HIS3; HPR1-TAP::URA3* | this study |
| *HPR1-TAP SYF1-HA Δcwc15* | BY4741 | *MATa; ura3Δ0; leu2Δ0; his3Δ1; met15Δ0; HPR1-TAP::URA3; SYF1-HA::HIS3; Δcwc15::kanMX4;* | this study |
| *HPR1-TAP SYF1-HA Δisy1* | BY4741 | *MATa; ura3Δ0; leu2Δ0; his3Δ1; met15Δ0; HPR1-TAP::URA3; SYF1-HA::HIS3; Δisy1::kanMX4* | this study |
| *HPR1-TAP SYF1-HA Δntc20* | BY4741 | *MATa; ura3Δ0; leu2Δ0; his3Δ1; met15Δ0; HPR1-TAP::HIS3; SYF1-HA::URA3; Δntc20::kanMX4;* | this study |
| *HPR1-TAP SYF1-HA Δsyf2* | BY4741 | *MATa; ura3Δ0; leu2Δ0; his3Δ1; met15Δ0; HPR1-TAP::URA3; SYF1-HA::HIS3; Δsyf2::kanMX4* | this study |
| *HPR1-TAP SYF1-HA Δsnt309* | BY4741 | *MATa; ura3Δ0; leu2Δ0; his3Δ1; met15Δ0; HPR1-TAP::URA3; SYF1-HA::HIS3; Δsnt309::kanMX4* | this study |
| *HPR1-TAP SYF1-HA Δcwc15 Δsyf2* | BY4741 | *MATa; ura3Δ0; leu2Δ0; his3Δ1; met15Δ0; HPR1-TAP::URA3; SYF1-HA::HIS3; Δsyf2::kanMX4; Δcwc15::LEU2;* | this study |
| *SYF1-TAP* | BY4741 | *MATa; ura3Δ0; leu2Δ0; his3Δ1; met15Δ0; SYF1-TAP::URA3;* | this study |
| *SYF1-TAP Δcwc15* | BY4741 | *MATa; ura3Δ0; leu2Δ0; his3Δ1; met15Δ0; SYF1-TAP::URA3; Δcwc15::kanMX4* | this study |
| *SYF1-TAP Δisy1* | BY4741 | *MATa; leu2Δ0; met15Δ0; his3Δ1; ura3Δ0, SYF1-TAP::URA3; Δisy1::kanMX4* | this study |
| *SYF1-TAP Δntc20* | BY4741 | *MATa; ura3Δ0; leu2Δ0; his3Δ1; met15Δ0; SYF1-TAP::URA3; Δntc20::kanMX4* | this study |
| *SYF1-TAP Δsyf2* | BY4741 | *MATa; ura3Δ0; leu2Δ0; his3Δ1; met15Δ0; SYF1-TAP::URA3; Δsyf2::kanMX4* | this study |

| | | | |
|---|---|---|---|
| *SYF1-TAP Δsnt309* | BY4741 | *MATa; leu2Δ0; met15Δ0; ura3Δ0 SYF1-TAP::URA3; Δsnt309*::kanMX4 | this study |
| *HPR1-TAP* | BY4741 | *MATa, his3Δ1; leu2Δ0; lys2Δ0; ura3Δ0; HPR1-TAP::URA3* | this study |
| *HPR1-TAP Δcwc15* | BY4741 | *MATa; ura3Δ0; leu2Δ0; his3Δ1; met15Δ0; HPR1-TAP::URA3; Δcwc15::kanMX4* | this study |
| *HPR1-TAP Δisy1* | BY4741 | *MATa; ura3Δ0; leu2Δ0; his3Δ1; met15Δ0; HPR1-TAP::URA3; Δisy1::kanMX4* | this study |
| *HPR1-TAP Δntc20* | BY4741 | *MATa; ura3Δ0; leu2Δ0; his3Δ1; met15Δ0; HPR1-TAP::URA3; Δntc20::kanMX4* | this study |
| *HPR1-TAP Δsyf2* | BY4741 | *MATa; ura3Δ0; leu2Δ0; his3Δ1; met15Δ0; Δsyf2::kanMX4; HPR1-TAP::URA3* | this study |
| *HPR1-TAP Δsnt309* | BY4741 | *MATa; ura3Δ0; leu2Δ0; his3Δ1; met15Δ0; HPR1-TAP::URA3; Δsnt309::kanMX4* | this study |
| *Δdst1* | W303 | *MATa; ura3-1; trp1-1; his3-11,15; leu2-3,112; ade2-1; can1-100; GAL+; Δdst1::HIS3* | this study |
| *Δcwc15* | W303 | *MATa; ura3-1; trp1-1; his3-11,15; leu2-3,112; ade2-1; can1-100; GAL+; Δcwc15::kanMX4* | this study |
| *Δisy1* | W303 | *MATa; ura3-1; trp1-1; his3-11,15; leu2-3,112; ade2-1; can1-100; GAL+; Δisy1::kanMX4* | this study |
| *Δntc20* | W303 | *MATa; ura3-1; trp1-1; his3-11,15; leu2-3,112; ade2-1; can1-100; GAL+; Δntc20::kanMX4* | this study |
| *Δsyf2* | W303 | *MATa; ura3-1; trp1-1; his3-11,15; leu2-3,112; ade2-1; can1-100; GAL+; Δsyf2::kanMX4* | this study |
| *Δsnt309* | W303 | *MATa; ura3-1; trp1-1; his3-11,15; leu2-3,112; ade2-1; can1-100; GAL+; Δsnt309*::kanMX4 | this study |
| *Δcwc15 Δdst1* | W303 | *MATa; ura3-1; trp1-1; his3-11,15; leu2-3,112; ade2-1; can1-100; GAL+; Δdst1::HIS3; Δcwc15::kanMX4* | this study |
| *Δisy1 Δdst1* | W303 | *MATa; ura3-1; trp1-1; his3-11,15; leu2-3,112; ade2-1; can1-100; GAL+; Δdst1::HIS3; Δisy1::kanMX4* | this study |
| *Δntc20 Δdst1* | W303 | *MATa; ura3-1; trp1-1; his3-11,15; leu2-3,112; ade2-1; can1-100; GAL+; Δdst1::HIS, Δntc20::kanMX4* | this study |
| *Δsyf2 Δdst1* | W303 | *MATa; ura3-1; trp1-1; his3-11,15; leu2-3,112; ade2-1; can1-100; GAL+; Δdst1::HIS3; Δsyf2::kanMX4* | this study |
| *Δsnt309 Δdst1* | W303 | *MATa; ura3-1; trp1-1; his3-11,15; leu2-3,112; ade2-1; can1-100; GAL+; Δdst1::HIS3; Δsnt309::kanMX4* | this study |
| *Δcwc15* | BY4741 | *MATa; his3Δ1; leu2Δ0; met15Δ0; ura3Δ0 Δcwc15::kanMX4* | Euroscarf |
| *Δisy1* | BY4741 | *MATa; his3Δ1; leu2Δ0; met15Δ0; ura3Δ0 Δisy1::kanMX4* | Euroscarf |
| *Δntc20* | BY4741 | *MATa; his3Δ1; leu2Δ0; met15Δ0; ura3Δ0 Δntc20::kanMX4* | Euroscarf |
| *Δsyf2* | BY4741 | *MATa; his3Δ1; leu2Δ0; met15Δ0; ura3Δ0 Δsyf2::kanMX4* | Euroscarf |
| *Δsnt309* | BY4741 | *MATa; his3Δ1; leu2Δ0; met15Δ0; ura3Δ0 Δsnt309::kanMX4* | Euroscarf |
| *TAP-HPR1 SYF1-HA* | BY4741 | *MATa; his3Δ1; leu2Δ0; met15Δ0; ura3Δ0; TAP-HPR1; SYF1-HA::HIS3* | this study |
| *TAP-HPR1 SYF1-HA Δcwc15* | BY4741 | *MATa; ura3Δ0; leu2Δ0; his3Δ1; met15Δ0; NTAP-HPR1; SYF1-HA::HIS3; Δcwc15::kanMX4* | this study |
| *TAP-HPR1 SYF1-HA Δsyf2* | BY4741 | *MATa; ura3Δ0; leu2Δ0; his3Δ1; met15Δ0; NTAP-HPR1; SYF1-HA::HIS3; Δsyf2::kanMX4* | this study |

| | | | |
|---|---|---|---|
| *TAP-HPR1 SYF1-HA Δcwc15 Δsyf2* | BY4741 | *MATa; ura3Δ0; leu2Δ0; his3Δ1; met15Δ0; NTAP-HPR1; SYF1-HA::HIS3; Δsyf2::kanMX4; Δcwc15::LEU24* | this study |
| *HPR1-TAP* | W303 | *MATa; ura3-1; trp1-1; his3-11,15; leu2-3,112; ade2-1; can1-100; GAL+; HPR1-TAP::TRP1* | this study |
| *HPR1-TAP Δcwc15 Δsyf2* | W303 | *MATa; ura3-1; trp1-1; his3-11,15; leu2-3,112; ade2-1; can1-100; GAL+; HPR1-TAP::TRP1; Δsyf2::kanMX4; Δcwc15::LEU2* | this study |

**Supplementary Table 2. Plasmids**

| **Plasmid** | **Description** | **Reference** |
|---|---|---|
| pBS1539 | for C-terminal TAP-tagging of a protein by genomic integration with the *URA3* marker | (3) |
| pBS1776 | for N-terminal TAP-tagging of a protein by genomic integration with the *LEU2* marker | (3) |
| pYM15 | for C-terminal HA-tagging with the *HIS3MX6* marker | Euroscarf |
| pRS316-*GAL10::ACT1* | *GAL10* promoter sequence in front of *ACT1* ORF in pRS316 | (4) |

**Supplementary Table 3. Oligonucleotides**

| **Primer** | **Sequence (5′–3′)** |
|---|---|
| YER-for | TGCGTACAAAAAGTGTCAAGAGATT |
| YER_rev | ATGCGCAAGAAGGTGCCTAT |
| PMA1-3'-for | CAGAGCTGCTGGTCCATTCTG |
| PMA1-3'-rev | GAAGACGGCACCAGCCAAT |
| DBP2-3'-for | CTTCACCGAACAAAACAAAGGTT |
| DBP2-3'-rev | TCGGGAGGAATATTTTGATTAGCT |
| ACT1-3'-for | TCAGAGCCCCAGAAGCTTTG |
| ACT1-3'-rev | TTGGTCAATACCGGCAGATTC |
| ILV5-3'-for | TGGTACCCAATCTTCAAGAATGC |
| ILV5-3'-rev | ACCGTTCTTGGTAGATTCGTACA |
| CCW12-3'-for | TGAAGCTCCAAAGAACACCACC |
| CCW12-3'-rev | AGCAGCAGCACCAGTGTAAG |
| RPL9B-3'-for | AGGACGAAATCGTCTTATCTGGT |
| RPL9B-3'-rev | CAGATTTGTTGCAAGTCAGCGG |
| RPL28-3'-for | TGGAAGCCAGTCTTGAACTTGG |
| RPL28-3'-rev | TTGGTCTCTCTTGTCTTCTGGGA |
| RPS14B-3'-for | AAGACCCCAGGACCAGGTG |
| RPS14B-3'-rev | GATACGGCCAATCCTCAAACCAG |
| ACT1-cy5 | Cy5- TGGATTGAGCTTCATCACCAACG |
| GAL10-cy5_96 | Cy5- ATTAGCTCTACCACAGTGTGTG |
| SCR1-cy5 | Cy5-TTTACGACGGAGGAAAGACG |
